# Repression of *Pdzrn3* is required for heart maturation and protects against heart failure

**DOI:** 10.1101/2020.07.29.226597

**Authors:** Mathieu Pernot, Béatrice Jaspard-vinassa, Alice Abelanet, Sebastien Rubin, Isabelle Forfar, Sylvie Jeanningros, Laura Cetran, Murielle Han-Yee Yu, Elise Balse, Stéphane Hatem, Pascale Dufourcq, Thierry Couffinhal, Cécile Duplàa

## Abstract

Heart failure is the final common stage of most cardiopathies. Cardiomyocytes connect with others via their extremities by intercalated disk protein complexes. This planar and directional organization of myocytes is crucial for mechanical coupling and anisotropic conduction of the electric signal in the heart. One of the hallmarks of heart failure is alterations in the contact sites between cardiomyocytes. Yet no factor on its own is known to coordinate cardiomyocyte polarized organization. We report enhanced levels of an ubiquitine ligase *Pdzrn3* in diseased hypertrophic human and mouse myocardium, which correlates with a loss of cardiomyocyte polarized elongation. We provide evidence that *Pdzrn3* has a causative role in heart failure. We found that cardiac *Pdzrn3* deficiency protected against heart failure while over expression of *Pdzrn3* in mouse cardiomyocytes during the first weeks of life, impaired postnatal cardiomyocyte maturation leading to premature death. Our results reveal a novel signaling pathway that controls a genetic program essential for heart maturation and maintenance of overall geometry, as well as the contractile function of cardiomyocytes, and implicates PDZRN3 as a potential therapeutic target for the prevention of human heart failure.

## Introduction

Heart failure develops gradually and is the final common stage of most cardiopathies. The hallmark of heart failure pathophysiology is the progressive nature of the disease, which evolves from a regional myocardial hypertrophy phase (concentric hypertrophy), viewed as an adaptive response of the heart to increased workload; to a decompensation, and dilation phase, resulting in heart failure (eccentric hypertrophy). At the cellular level, alteration of the organization and molecular composition of contact sites between cardiomyocytes, the intercalated discs (IDs) was related to be associated with heart failure (Ehler et al., 2001; Ferreira-Cornwell et al., 2002; Phillips et al., 2007).

Cardiomyocytes can be seen as highly polarized cells with specialized structures present only between the ends of each abutted cell, the IDs. These regions provide cell-to-cell mechanical connections and mediate electrochemical communication(E. Balse et al., 2012). During the embryonic stage, myocardial cells start to be locally coordinated and aligned in a planar bias(Le Garrec et al., 2013). Initially spherical, cardiomyocytes gradually elongate during foetal and postnatal development, to adopt a rod shape. Most components of ID are initially expressed all around the foetal cardiomyocyte membrane and, after birth, following the elongation process, they progressively reorganize from the lateral cell membrane to the ends of each cell. The remodeling of these specialized organized regions of the plasma membrane is only achieved after birth. Thus, a deeper understanding of the molecular mechanisms governing postnatal establishment and maintenance of these IDs, and alterations during the development of heart failure is a critical step toward developing novel therapeutic strategies to prevent these pathologies. Desmosome molecules are major architectural components of the IDs in cardiac muscle, building blocks which are intermixed with components of adherens and gap junctions (Estigoy et al., 2009). Fascia adherens junctions like N-cadherins (N-Cad) form complexes with the α-, β-, and γ-catenin proteins at the adherens junctions and facilitate calcium-dependent cell adhesion between adjacent cardiac myocytes and attachment sites for myofibrils (Mezzano et al., 2014). Connexin43 (Cx43) has been identified as the principal component of gap junctions in ventricular myocytes(Delmar & Liang, 2012). The connexins provide a low resistance portal for the transmission of electrical impulses and chemicals between cells.

Planar cell polarity (PCP) signalling is a main pathway involved in tissue patterning, instructing cell polarity and cell polar organization within a tissue(Yang & Mlodzik, 2015). It ensures a coordinated planar polarization between contacting cells. Initially identified in Drosophila, the PCP pathway is well conserved in mammalian species. PCP signalling involves a multiprotein complex that associates at the cell membrane, including Frizzled, Dishevelled (Dvl), Vangl (Vangl), and Scribble (Scrib). PCP pathways are important in polarized cell migration and organ morphogenesis through activation of cytoskeletal pathways, such as those involving the small GTPases RhoA and Rac, protein kinase C, and Jun N-terminal kinase. It is also involved in regulating cardiogenesis in vertebrates (Pandur et al., 2002). Genetic mouse studies have demonstrated that PCP components such as Wnt11, Vangl2, Scrib, Dvl2 and Rac1 regulate cardiomyocyte polarity and embryonic heart development(Henderson et al., 2001; Leung et al., 2014; Nagy et al., 2010). Deletion of Wnt-11 and Vangl2 reveals its critical role in cell adhesion required for organization of cardiomyocytes in the developing ventricular wall (Henderson et al., 2001; Nagy et al., 2010). However, the role of PCP in postnatal myocyte shape maturation and polarized ID reorganization has not yet been investigated.

We have previously shown that PDZRN3, an ubiquitine ligase E3 expressed in various tissues including the heart, mediates a branch of the PCP pathway involved in vascular morphogenesis and in the stabilization of endothelial cell/cell junctions(R. N. Sewduth et al., 2014; Raj N. Sewduth et al., 2017). It has also been reported to be required for myocyte(Ko et al., 2006) and osteoblast differentiation(Honda et al., 2010). In the present study, we demonstrate, using loss and gain of function genetic approaches, that *Pdzrn3* is a critical regulator of cardiomyocyte maturation and stabilization of IDs and plays a major role in the transition from concentric to eccentric hypertrophy toward heart failure. Similar features are reported in human heart samples. We also reveal that PDZRN3 regulates Wnt signalling in cardiomyocytes. This study identifies PDZRN3 as a novel signalling regulator of polarized myocyte organization and opens this pathway up to the development of novel therapeutic strategies for preventing heart failure pathologies.

## Results

### Loss of *Pdzrn3* protects the transitions of the heart from compensated to decompensated cardiac hypertrophy

*Pdzrn3* was initially identified by two-hybrid screening of a mouse 12.5 embryonic day (E12.5) heart cDNA library(R. N. Sewduth et al., 2014). Reactivation of fetal gene programs have been observed in pathological processes underlying maladaptive changes in cardiac hypertrophy(Heineke & Molkentin, 2006; Hoshijima & Chien, 2002), therefore we asked whether *Pdzrn3* gene expression was related to myocardial hypertrophic adaptation.

We examined *Pdzrn3* expression in human heart samples from patients with adaptative or decompensated hypertrophic cardiomyopathy (respectively control-HCM or HF-HCM), and with decompensated dilated cardiomyopathy (HF-DCM) and in two mouse models of pressure overload. Using quantitative PCR analysis, we found that *Pdzrn3* mRNA level was significantly increased in tissue samples retrieved from patients with adaptative or decompensated HCM (control-HCM) compared to HF-DMC samples, with HF-HCM having higher levels than HF-DMC samples. (Fig. 1a). In mice we found that, pressure-overload stress of the heart by Angiotensin-II (Ang II) infusion or transverse constriction (TAC) surgery, caused cardiac hypertrophy and lead to heart failure (decompensation phase). We found an increased abundance of transcripts for *Pdzrn3* associated with heart failure in both mouse models 4 weeks post Ang II treatment (Fig 1b) and 16 weeks post TAC surgery (Fig 1c).

**Figure 1.**
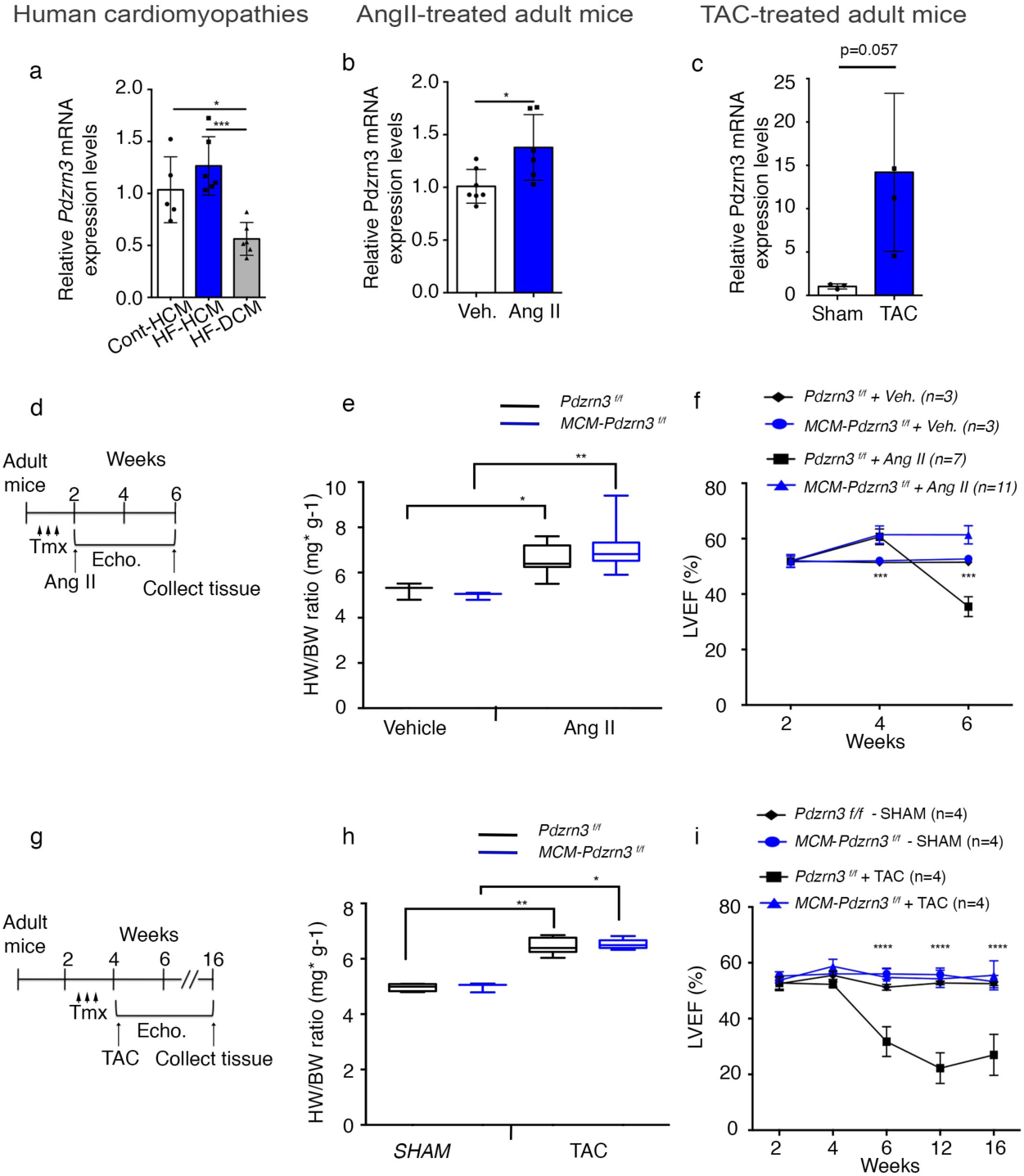
Loss of Pdzrn3 protects the transition of the heart from compensated to decompensated cardiac hypertrophy. a. Quantitative real-time PCR analysis of Pdzrn3 transcript abundance in hearts retrieved from patients with non-decompensated (control HCM), decompensated HCM (HF-HCM), and from decompensated primitive or ischemic dilated cardiomyopathy (HF-DCM). mRNA levels were normalized to GAPDH and are expressed as relative expression over levels in the control-HCM group (n=5 in control-HCM and n=6 in other groups). b. Quantitative real-time PCR analysis of Pdzrn3 transcript abundance in hearts retrieved from wild type mice after treatment with saline (Veh.) or Ang II for 4 weeks. mRNA levels were normalized to cyclophiline and are expressed as relative expression over levels in the control Veh. treated group (n=3-7 per group). c. Quantitative real-time PCR analysis of Pdzrn3 transcript abundance in hearts retrieved from wild type mice after TAC surgery. (sham group, n=3, TAC group, n=4). mRNA levels were normalized to cyclophiline and are expressed as relative expression over levels in the control Sham group. d. Schematic study plan showing timeline of tamoxifen administration (Tmx), Ang II pump implantation, echocardiography (Echo.) follow up and tissue collection. e. Ratio of heart weight (HW) to body weight (BW). f. Quantification of left ventricular ejection fraction (LVEF) in tamoxifen treated *Pdzrn3*^*f/f*^ and MCM-Pdzrn3 mice following Ang II treatment. (vehicle treated groups, n=3; Ang II-treated groups, *Pdzrn3*^*f/f*^ n=7, MCM-Pdzrn3, n=11), g. Schematic study plan showing timeline of tamoxifen administration (Tmx), TAC surgery, echocardiography (Echo.) follow up and tissue collection. h. Ratio of heart weight (HW) to body weight (BW). i. LVEF quantification in tamoxifen treated *Pdzrn3*^*f/f*^ and MCM-*Pdzrn3* mice following TAC surgery (SHAM groups, n=3 ; TAC-treated groups, *Pdzrn3*^*f/f*^ n=7, MCM-Pdzrn3, n=5). Mean ± s.e.m. *P<0.05, ** P<0.005, ***, P<0.001 by one way ANOVA (a) and non parametric *t*’test (b, c, e, h), repeated-measures two way ANOVA with tukey’s test (f, i).

We next investigated whether *Pdzrn3* is required during the development of cardiac mouse hypertrophy. We crossed mice bearing a *Pdzrn3* flox allele (*Pdzrn3*^*f/f*^) with transgenic mice (MerCreMer) that express Cre recombinase in a tamoxifen-inducible and cardiomyocyte-specific manner (*24*) (Suppl fig. S1a). The resulting MCM-*Pdzrn3*^*f/f*^ mice were indistinguishable in appearance from age-matched control *Pdzrn3*^*f/f*^ littermates. In 8 week-old MCM-*Pdzrn3*^*f/f*^ mice, treated with tamoxifen for 3 consecutive days, efficient loss of *Pdzrn3* transcripts was observed (Suppl fig. S1b). There was no alteration in left ventricular ejection fraction (LVEF) and other cardiac parameters associated with *Pdzrn3* deletion as analysed by echocardiography over 6 months (data non shown).

Interestingly, we found that tamoxifen-treated MCM-*Pdzrn3*^*f/f*^ mice are resistant to Ang II-induced cardiac maladaptive hypertrophy (Fig. 1d). Ang II infusion induced in both *Pdzrn3*^*f/f*^ and MCM-*Pdzrn3*^*f/f*^ mice development of a left ventricular hypertrophy as showed by an increase in heart-to-body weight ratios compared to both vehicle treated *Pdzrn3*^*f/f*^ or MCM-*Pdzrn3*^*f/f*^ mice (Fig. 1e). Two weeks after Ang II infusion, both *Pdzrn3*^*f/f*^ and MCM-*Pdzrn3*^*f/f*^ mice showed a significant increase of LVEF (Fig. 1f). One month after Ang II treatment, *Pdzrn3*^*f/f*^ mice showed an eccentric heart hypertrophy and severe contractile dysfunction, whereas MCM-*Pdzrn3*^*f/f*^ mice maintained their LVEF at the same level as seen at earlier time points (Fig.1f).

Similar results were obtained using a model of TAC surgery (Fig. 1g). Tamoxifen-treated *Pdzrn3*^*f/f*^ and MCM-*Pdzrn3*^*f/f*^ mice were subjected to TAC pressure overload for 16 weeks. Four weeks after the onset of surgery, *Pdzrn3*^*f/f*^ and MCM-*Pdzrn3*^*f/f*^ mice developed significant left ventricular concentric hypertrophy (Fig. 1h). During the follow-up, only *Pdzrn3*^*f/f*^ mice developed a significant alteration in LVEF as early as 6 weeks after undergoing the surgical TAC procedure. MCM-*Pdzrn3*^*f/f*^ mice, showed complete protection against degradation of heart function. LVEF in MCM-*Pdzrn3*^*f/f*^ mice was maintained at normal levels up to 4 months after TAC surgery (Fig. 1i).

All together these data indicate that in cardiomyocytes, *Pdzrn3* is essential in the transition from concentric to eccentric hypertrophy and subsequent heart failure in models of induced cardiac hypertrophy.

### Myocardial *Pdzrn3* reactivation induces cardiac eccentric hypertrophy

To further understand the function of *Pdzrn3* induction in cardiac disease, we analysed myocardial expression patterns of PDZRN3. Western blot analysis confirmed that PDZRN3 is expressed in the embryonic mouse heart (E 11.5 to 15.5), but its expression dropped significantly postnatally from day (d) 0.5 to 14 d, becoming faintly expressed in the adult mouse heart (2 months). This kinetic corresponds with heart maturation (Suppl fig. 2a).

To address its role in cardiac maturation, we employed a gain-of-function approach to overexpress Pdzrn3 in postnatal myocardium. Mice overexpressing Pdzrn3 (*Pdzrn3* OE) were generated by crossing reporter mice, which harbor a bidirectional *Tet*-promoter cassette with genes for PDZRN3-V5 and β galactosidase, with *αMHC-tTA* mice to allow for specific expression of *Pdzrn3* in cardiomyocytes at birth. (Suppl Fig. 2b). We used Western blot to examine the expression pattern of ectopic PDZRN3 related to endogenous PDZRN3 levels during postnatal maturation. Ectopic PDZRN3 could be detected after birth at 0.5 d, and peaked between 4 and 7 d by approximately 2- and 3-fold, respectively. After 14 d, ectopic PDZRN3 decreased to levels seen endogenously at 0.5 d, and maintained low levels of ectopic PDZRN3 expression into adulthood (Suppl Fig. 2c-d).

We observed that LVEF was progressively impaired as early as 4 weeks, in *Pdzrn3* OE mice compared with control mice, resulting in nearly 100% mortality at 28 weeks of age (Fig. 2a-b). Surface electrocardiograms (ECGs) performed on 3-10-week-old *Pdzrn3* OE mice and their control littermates showed no differences in heart rate or RR, QRS and QTc intervals. However, *Pdzrn3* OE mice exhibited a prolongation of the PR interval as early as 3-week-old (46+/- 4 vs 35+/-2 ms, p<0.01) (Suppl Fig. 3), and this first-degree heart block remained constant until death. Approximately 20% of the mice displayed auriculoventricular blocks (Mobitz I and II) but no third-degree AV block. No ventricular arrhythmias were recorded. Cardiac overexpression of *Pdzrn3* led to an eccentric hypertrophic phenotype (Fig. 2c). At 6 weeks, *Pdzrn3* OE mice exhibited an increase in the internal cross-sectional area of the left ventricle compared to their control littermates, an increased in the heart-to-body weight ratio, and an expected increase in mRNA expression of molecular markers for cardiac remodelling, including atrial natriuretic factor (*Nppa*) and brain natriuretic peptide (*Nppb*) (Fig. 2d-f). As early as 6 weeks, histological analysis revealed an increase in fibrosis and inflammation in the hearts of *Pdzrn3* OE mice (Fig. 2g). Furthermore, glycogen accumulation was confirmed by the presence of periodic schiff positive materials in 4 week old mice (Fig. 2g).We then monitored and quantified an increase in glycogen storage *in situ* by the Fourier Transform Infrared Spectroscopy (FTIR) method in heart tissue sections of mutant mice (Fig. 2h) at two and four weeks of age, suggesting that precocious overexpression of *Pdzrn3*, during the first weeks of life, can affect cardiomyocyte metabolism which may cause or precede the development of heart failure.

**Figure 2.**
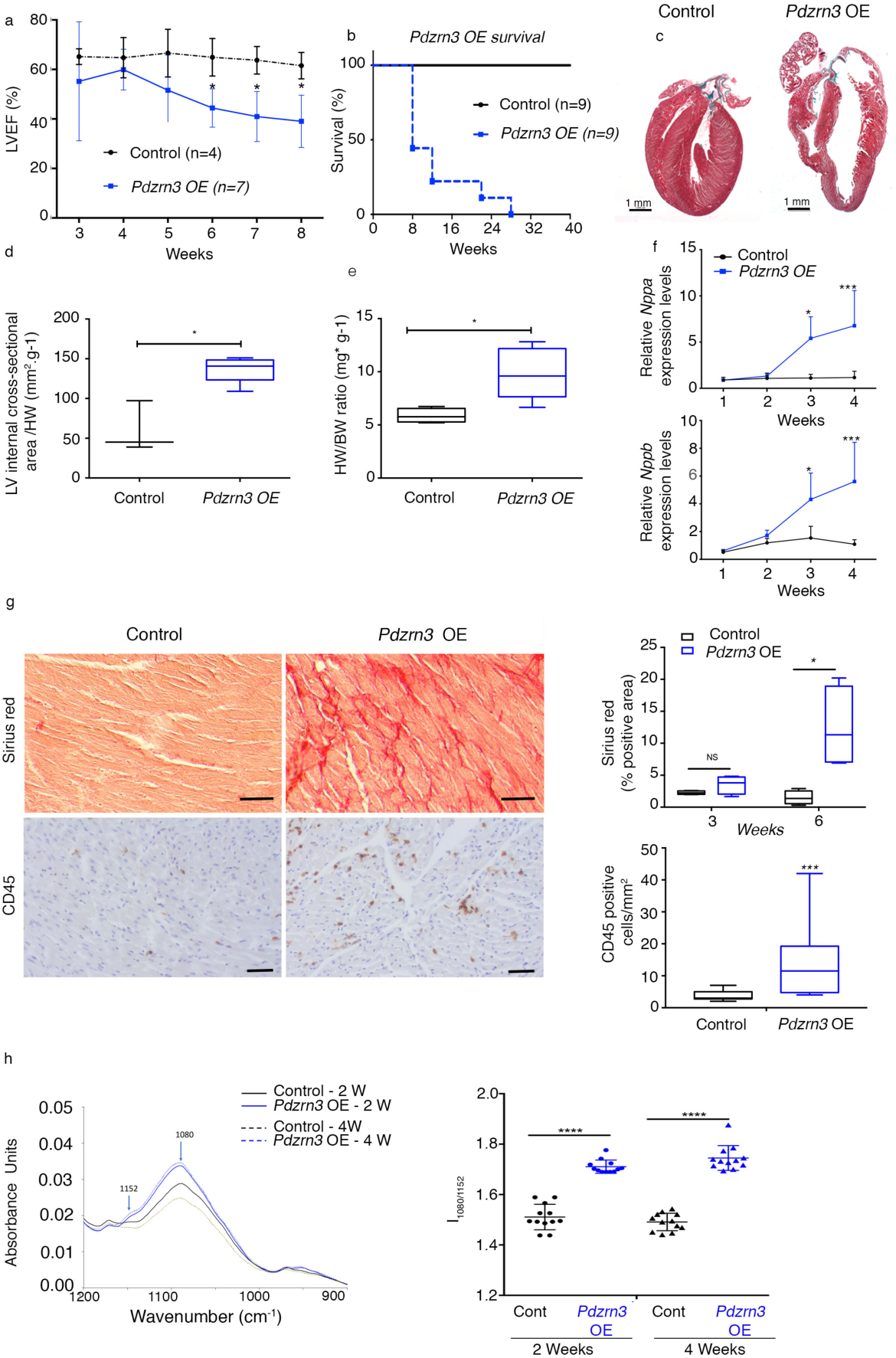
Myocardial reactivation of *Pdzrn3* induces cardiac eccentric hypertrophy. a. LVEF was quantified from 3 to 8 weeks of age (control, n= 4; *Pdzrn3* OE, n=7). b. Kaplan-Meier curve showing survival rate of control and *Pdzrn3* OE mice from birth until 40 weeks of age (control and *Pdzrn3* OE groups, n= 9). c. Representative images of hematoxylin and eosin (H&E) stained sections of control and *Pdzrn3* OE hearts at 6 weeks, demonstrate that Pdzrn3 OE mice have enlarged hearts. d. Increased LV internal cross section (control, n=3; *Pdzrn3* OE, n= 5). e. Increased heart weight (HW) to body weight (BW) ratio (control, n=5; *Pdzrn3* OE, n= 5). f. Increase of the mRNA levels of atrial (Nppa) and brain (Nppb) natriuretic peptide in *Pdzrn3* OE versus control mice over time. mRNA levels were normalized to cyclophiline and are expressed as relative expression/ fold increase over levels found in control mice at 1 week (n=3-7 mice per group) g. In hearts from *Pdzrn3* OE mice, quantification of total collagen deposition on picrosirius red staining revealing an increase in fibrosis and CD45 immunolabeling, and an increase of inflammatory cells at 6 and 4 weeks of age, respectively (n=4 mice per group) (the scale bars represent 50 μm). h. FTIR spectra after pre-processing and classification (n=6 spectra per condition) in the 900 cm^-1^ to 1200 cm^-1^ spectral region obtained heart tissue cryo-sections retrieved from the left ventricle from control and *Pdzrn3* OE mice (n=3 per group at 2 and 4 weeks). Arrows indicate the wavenumbers studied. Changes in the heart glycogen illustrated with the absorbance ratio 1080 cm^-1^/1152 cm^-1^. All the data are represented as mean ± sem, with n=3 mice. Mean ± s.e.m. *P<0.05, ** P<0.005, ***, P<0.001 by repeated-measures two way ANOVA with post-hoc sidak’s test (a, f); Kaplan-Meier non parametric regression analysis and the log-rank test (b) and unpaired *t*-test (d, e, g).

### Pdzrn3 signaling is required for cardiomyocyte polarized elongation along the development of heart disease in adult and postnatally

Cardiac hypertrophy to heart failure transition was reported to activate cardiomyocyte gene programs that orchestrate morphological phenotypes (Nomura et al., 2018). Here we hypothesized that the loss of *Pdzrn3* may contribute to the mechanism of protecting the heart from transitions from a compensated to a decompensated cardiac hypertrophy in promoting cardiomyocyte elongation. Cross–sectional area of cardiomyocytes was measured using FITC– wheat germ agglutinin–stained sections and the sphericity index was followed as an index of changes in myocyte shape (Fig. 3a). We measured this index in human cardiac tissues, and reported an increase in the cardiomyocyte sphericity index in HF-HCM compared to HF-DCM heart tissues (Fig. 3b). In adult mice treated with AngII for 4 weeks, the sphericity index was significantly increased in *Pdzrn3*^*f/f*^ compared to MCM-*Pdzrn3*^*f/f*^ mice while no significant change in the sphericity index was observed in vehicle-treated *Pdzrn3*^*f/f*^ and MCM-*Pdzrn3*^*f/f*^ mice (Fig. 3c). Finally, as cardiomyocyte elongation and alignment starting from the perinatal stage is a critical step in cardiomyocyte maturation, we test the hypothesis that maintaining PDZRN3 signaling during this postnatal period in mice may perturbate cardiomyocyte maturation. We report that ventricular cardiomyocytes underwent a specific lengthening 2 to 4 weeks after birth, consistent with cardiomyocyte maturation and that forced ectopic *Pdzrn3* expression impaired shape changes in cardiomyocytes which remained in a round shape (Fig. 3d). Ultrastructural analysis of hearts from *Pdzrn3* OE mice confirmed this analysis and revealed that cardiomyocyte loss their planar polarized organization and acquired a round phenotype at 8 weeks of age (Fig. 3e).

**Figure 3.**
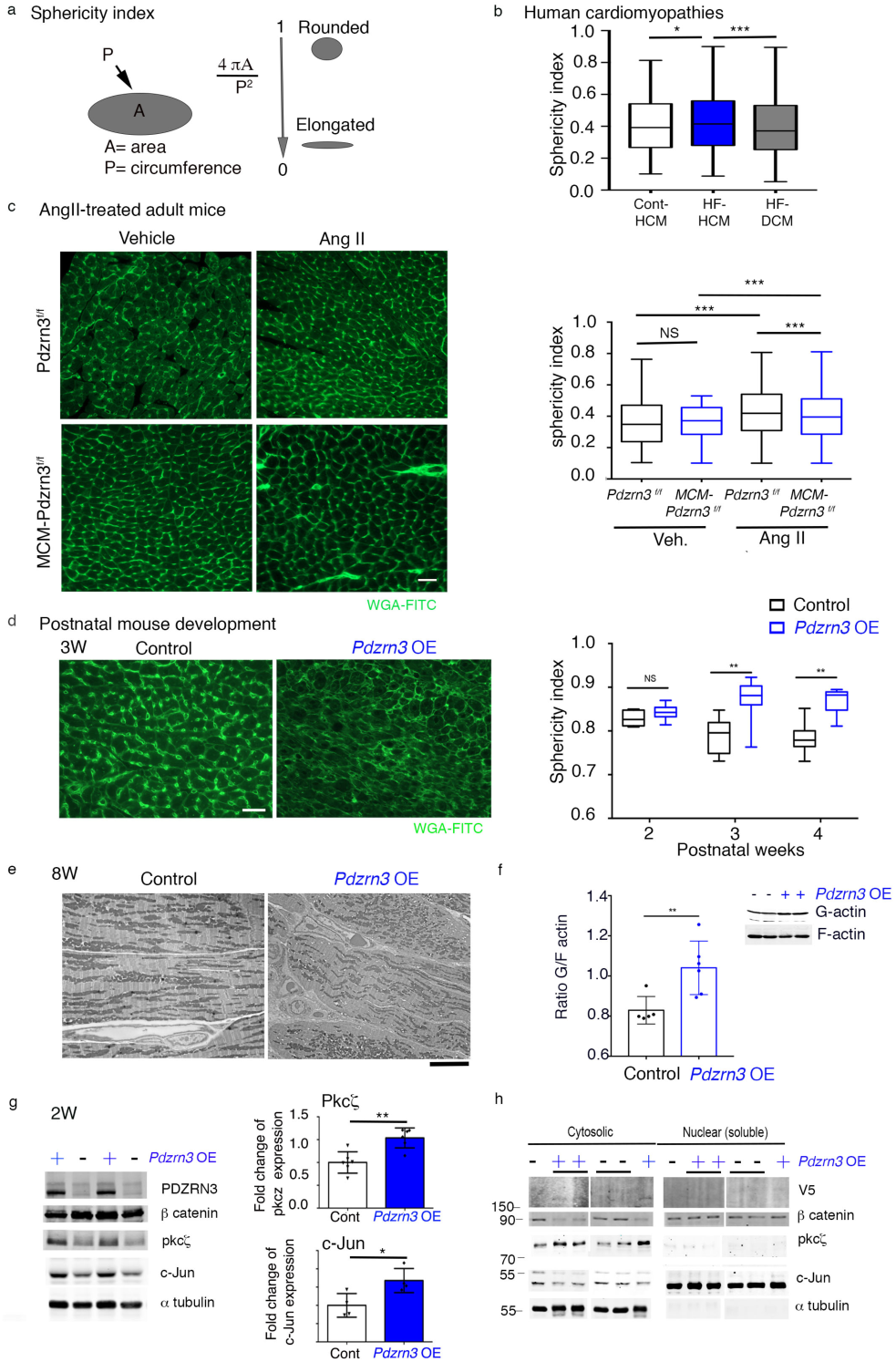
Cardiac specific overexpression of *Pdzrn3* impairs cardiomyocyte elongation. a. The sphericity index was determined by measuring the area (A) and the circumference (P) of myocytes. The ratio of an “ideal” round cell is close to 1, whereas that of an elongated cell is closer to 0. b. Quantification of the cardiomyocyte sphericity index in heart sections from patients with non-decompensated (control HCM), decompensated HCM (HF-HCM), and from decompensated primitive or ischemic dilated cardiomyopathy (HF-DCM). (n=3 in control-HCM, n=4 in HF-HCM, n=6 in HF-DCM). Mean ± s.e.m. *P<0.05, ** P<0.005 by one way ANOVA with Dunn’s test (b, d, e). c. Quantification of the cardiomyocyte sphericity index in tamoxifen treated *Pdzrn3*^*f/f*^ and MCM-Pdzrn3 mice following Ang II treatment. (vehicle treated groups, n=3; in Ang II-treated groups, *Pdzrn3*^*f/f*^ n=3, MCM-Pdzrn3, n=6). d. Representative images of control and *Pdzrn3* OE heart sections stained with wheat germ agglutinin (WGA)-FITC (scale bars represent 20 μm) at 3W. Quantification of myocyte cross-sectional sphericity of control and *Pdzrn3* OE heart sections. Data are represented as mean ± sem. Significance vs control **, P<0.01 by one way ANOVA plus bonferroni test. (control vs *Pdzrn3* OE group, n= 4 at 2 weeks, n=3 at 3 weeks and n=5 at 4 weeks). e. Representative electron micrographs showing morphological disorganization of cardiomyocytes of control and *Pdzrn3* OE heart at 8 weeks of age (scale bars represent 1 μm) f. Quantitative F-actin/G-actin ratios in heart lysates from 2 week old mice were measured with actin polymerization *in vivo* assay kit. Western blot analysis of G-actin (G) and F-actin (F) fraction from WT and Pdzrn3 OE heart extracts were probed with anti-actin antibody. Ratios of F-actin/G actin were determined from the blots by optical density measurements. \*\**p* < 0.001, by unpaired *t*-test. g. Western blot analysis of indicated proteins in heart tissues retrieved from control and *Pdzrn3* OE mice at 2 weeks (W) of age. Relative expression of Pkc *ζ* and c-Jun from Western blots. Significance vs control **, P<0.01; ***, P<0.001 by unpaired *t*-test (at 1 week, n=4 mice per group; at 2 weeks, n=6 mice per group). h. Western blot analysis of indicated proteins in heart tissue retrieved from control and *Pdzrn3* OE mice after tissue fractionation to isolate cytosolic, membrane and nuclear (soluble) fractions. (n=3 mice per group).

Pdzrn3 was known for its involvement in cytoskeletal rearrangement(Raj N. Sewduth et al., 2017). Because actin cytoskeleton is involved in morphogenetic process as elongation (Chhabra & Higgs, 2007), we analyzed the relationship between actin and Pdzrn3. The G/F-actin ratio significantly increased in the Pdzrn3 OE condition, suggesting an impairment or reduction in actin polymerization state in cardiomyocyte where Pdzrn3 levels were increased (Fig. 3f). To further investigate the molecular signature correlated with impair cardiomyocyte postnatal maturation and planar polarity, we examined previously shown mediators of the PCP pathway as PKCζ and c-Jun. We found that PKCζ and c-Jun protein levels were significantly increased at 14 d in heart lysates from *Pdzrn3* OE mice (Fig. 3g). A protein fractionation strategy was then employed to separate and enrich the various cellular structures from *Pdzrn3* OE mice and control littermate tissue samples at 14 d (Fig. 3h). The amount of PKCζ protein was increased in the cytosol fraction while c-Jun was translocated in the nucleus. Total β catenin protein level was decreased in the cytosol fraction but not at the membrane. All together, these data suggest that PDZRN3 signalling in cardiomyocytes, favors c-Jun and PKCζ activation, molecular actors of the Wnt non-canonical pathway, while it impairs β catenin canonical Wnt signalling.

### Myocardial *Pdzrn3* reactivation alters postnatal cardiomyocyte maturation

To identify protein biomarkers specific for impaired postnatal cardiomyocyte polarization under *Pdzrn3* ectopic expression, a mass spectrometry (MS) quantitative analysis was performed on heart lysates from *Pdzrn3* OE mice and control littermate tissue samples at 14 d. 4,363 proteins were identified and among these, 39 proteins were differentially regulated, which includes 23 up-regulated and 16 down-regulated proteins (with p value <0.05 set as significant) (Suppl Table 1). Gene-ontology (GO) analyses were then carried out and we found that several proteins are functionally associated with the maintain of cell junction and cell leading edge (Suppl Table 2). Consistent with this, by electron microscopy we show a massive regression of junctions at the longitudinal cell-edges between cardiomyocytes at 2 months old *Pdzrn3* OE mice (Fig. 4a).

**Figure 4.**
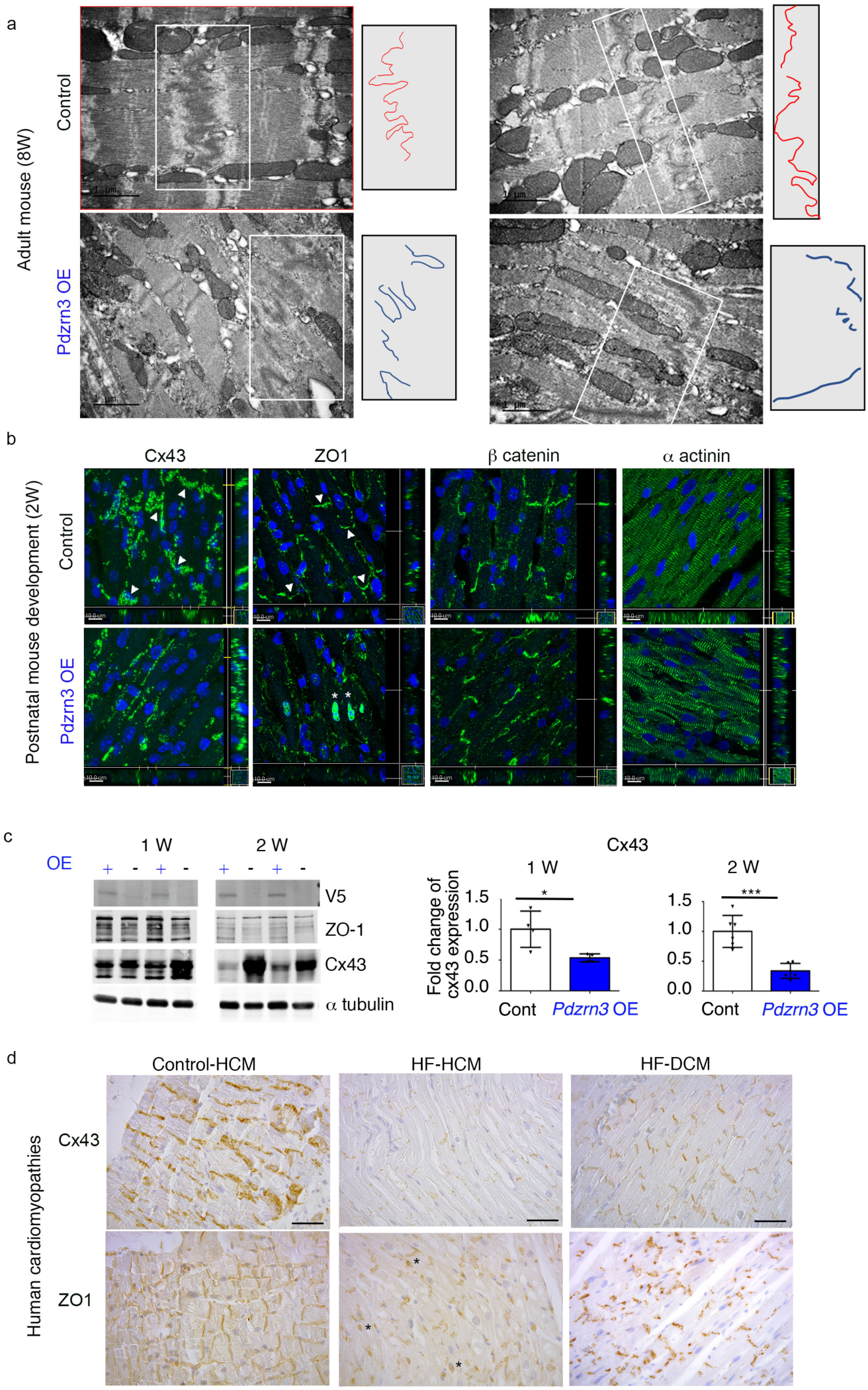
Myocardial reactivation of Pdzrn3 alters post natal cardiomyocyte maturation. **a**. Electron micrographs of part of an intercalated disc between two cardiac of control and *Pdzrn3* OE heart at 8 weeks of age (scale bars represent 2 μm) and in right part, schema of ID distribution. **b**. Immunolabeling of Cx43, ZO1, β catenin and *α* actinin in hearts from control and *Pdzrn3* OE mice at 2 weeks. Scale bars represent 10 μm. **c**. Western blot analysis of indicated proteins in heart tissues retrieved from control and *Pdzrn3* OE mice at 1 and 2 weeks (W) of age. Relative expression of Cx43, from Western blots. Significance vs control *, P<0.05 ; ***, P<0.001 by unpaired *t*-test (at 1 week, n=4 mice per group; at 2 weeks, n=6 mice per group). **d**. Representative immunolabeling of Cx43 and ZO-1 in tissues from patients with non-decompensated (control HCM), decompensated HCM (HF-HCM), and from decompensated primitive or ischemic dilated cardiomyopathy (HF-DCM). Scale bars represent 50 μm.

These results prompted us to hypothesize that forced *Pdzrn3* expression may impair localization of cell-cell junctional components during the initial myocyte maturation steps in mice. Henceforth, analysis of proteins that localize to ID were prioritized. First, we focused on Cx43 expression as Cx43 (*Gja1* gene) sorted out as a specific downregulated biomarker at 2 weeks in *Pdzrn3* OE mouse mutant hearts (Suppl Table 1). Immunohistological staining confirms MS analysis; the expression of Cx43 decreased as early as 1 week and was dramatically reduced at 2 weeks in heart sections of *Pdzrn3* OE mice. At 2 weeks, Cx43 expression is still detected in lateral membrane but not in ID as compared to control littermates (Fig. 4b). Western blot semiquantitative analysis confirmed the significant decrease of Cx43 expression at 7 d and 14 d (Fig. 4c).

ZO-1 re-localization from the lateral cell membrane toward the IDs at 14 d was impaired in *Pdzrn3* OE mice compared to control littermates (Fig. 4b). We found also an unexpected translocation of ZO1 in the nucleus which may be correlated with a decrease of maturity of cell/cell contact (Gottardi et al., 1996). At that time, we did not observe modification in expression and localization of other junctional proteins, including β catenin and N cadherin in mutant hearts (Fig. 4b and suppl Fig 4).

Based on our findings that Pdzrn3 was upregulated in human HF-HCM hearts, we next investigated whether the development of heart failure in humans might be associated with the regulation of the expression of the biomarker Cx43 and cellular localization of ZO-1. In tissues from patients suffering of HF-HCM, we found large areas in which Cx43 was completely depleted and ZO-1 was re-localized around the lateral membrane of cardiomyocytes and also translocate in the nucleus. This specific feature was not found in HF-DCM or control-HCM tissues. (Fig. 4d).

### Narrow postnatal time window is required for cardiac cell differentiation

Having shown that PDZRN3 is a critical trigger in controlling postnatal cardiomyocyte maturation, and in light of previous reports that cardiomyocyte maturation must be completed within a short time window(Porrello & Olson, 2014), we conducted experiments to assess whether, transgenic *Pdzrn3* overexpression within a restricted postnatal time period, is sufficient to alter cardiomyocyte maturation and to induce an eccentric hypertrophic response in adult mice. Doxycycline, a stable analogue of tetracycline, was delivered to mice in the drinking water after weaning, to block *Pdzrn3* expression. Mice were administrated with doxycycline either from 0 d (birth) (Fig. 5 a) or from 7 d (Fig. 5 b) and sacrificed at 14 d for histological analysis. In both protocols, Western blot analysis confirmed that doxycycline treatment impaired ectopic PDZRN3 expression. As expected, repression of ectopic *Pdzrn3* expression from birth abrogated the decrease of Cx43 expression and the mis-localization of ZO-1 (Fig. 5 c-d) while overexpression of *Pdzrn3* only during the first week (7-14 d doxycycline treatment) was sufficient to significantly decrease Cx43 expression and maintain ZO-1 expression in lateral cardiomyocyte membranes (Fig. 5 e-f). This data suggests that ectopic PDZRN3 signalling in the first week of life represses Cx43 expression and ZO1 re-localization.

**Figure 5.**
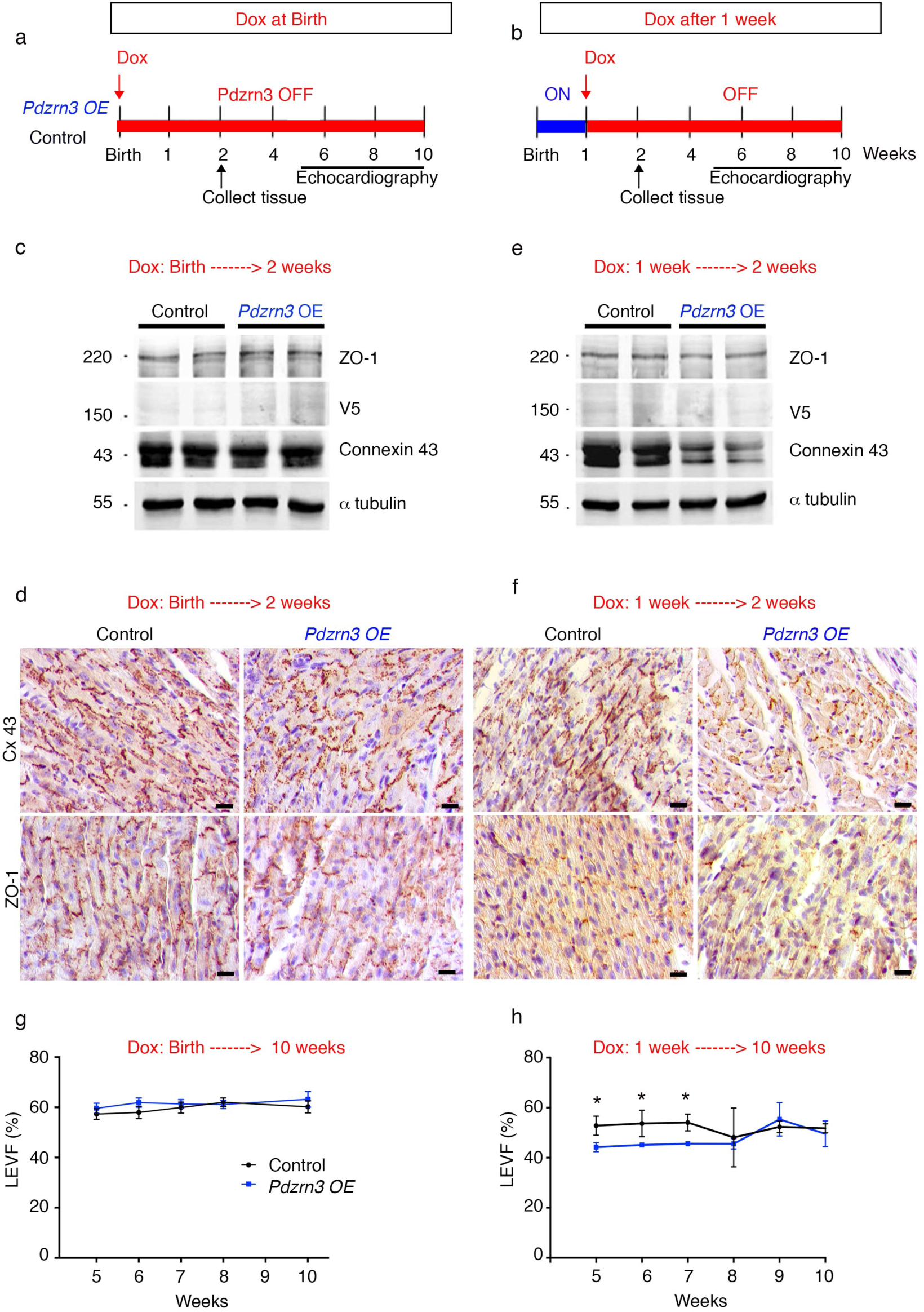
Postnatal time window for cardiac cell differentiation. a.and b. Schematic study timeline of doxycline treatment administration (Dox), tissue collection and echocardiography follow up. c. and e. Western blot analysis of indicated proteins in heart tissues from control and Pdzrn3 OE mice. d. and f. Immunolabeling of Cx43 and ZO1 in hearts from control and *Pdzrn3* OE mice following doxycycline treatment. Scale bars represent 50 μm. g. and h. LVEF quantification under doxycycline treatment in control and Pdzrn3 OE mice in d protocol, respectively n=6 vs 8; in h protocol, respectively n=6 vs 4 mice.

To examine long-term effects of restricted ectopic *Pdzrn3* expression during the first week of life, cardiac function was analysed for up to 4 months under doxycycline treatment. When *Pdzrn3* postnatal expression was repressed at birth, no signs of cardiomyopathy were detected; cardiac structure, dimension and LVEF were all normal (Fig. 5g). *Pdzrn3* OE mutant mice treated with doxycycline beginning at 7 d exhibited impaired LVEF at 8 weeks; interestingly, this alteration did not persist after 10 weeks (Fig. 5h).

All together, these data suggest that PDZRN3 is a critical master regulator of cardiomyocyte postnatal maturation and cardiac morphogenesis.

### Loss of *Pdzrn3* increases Desmoglein accumulation at cell membrane

Alteration in adherent junction are reported to precede Cx43 gap junction changes (Elise Balse et al., 2012; Yoshida et al., 2011). We then hypothesized that *Pdzrn3* may impact on the maintain of junctional complex of cardiac muscle. The cadherin-like adhesion molecule Desmoglein (Dsg) is crucial for cardiomyocyte cohesion and is dynamically localized in ID during the first 2 weeks after birth and functions to maintain tissue integrity (Krusche et al., 2011; Schlipp et al., 2014). MS quantitative analysis did not show significant decrease of overall Dsg2 levels at 2 weeks in heart lysate from mutant compared to control hearts. We then performed subcellular fractionation from heart and revealed that Dsg2 protein levels were decreased in plasma membrane fraction of heart lysates from *Pdzrn3* OE mice compared to control mice at 2 weeks by Western blot analysis (Fig 6a). Using immunofluorescence analysis, we confirmed this morphological phenotype on neonatal cardiomyocytes isolated from *Pdzrn3* OE mice vs control littermate mice with a decrease expression of Dsg2 and Cx43 at cell/cell junctions and a translocation of ZO1 in the nucleus (Fig 6b). *In vitro* studies were then conducted to analyse whether Dsg2 accumulation at cell membrane is regulated by *Pdzrn3*. We found that repressing *Pdzrn3* expression in HeLa cells using siRNA increases accumulation of Dsg2 at cell/cell contact by immunofluorescent assays (Fig 6c). Many components of cell junctions are insoluble in detergents as triton X-100 (Tx-100) due to their association with the cortical actin at the cell periphery. Dsg2 expression level increases in Triton X100-insoluble flotillin enriched membrane fraction of *Pdzrn3* depleted cells suggesting that PDZRN3 is a key determinant of Dsg2’s localization at cell-cell junctions (Fig 6d). Overall, these findings confirm that PDZRN3-induced signaling controls the dynamic junction complex assembly at cell periphery.

**Figure 6.**
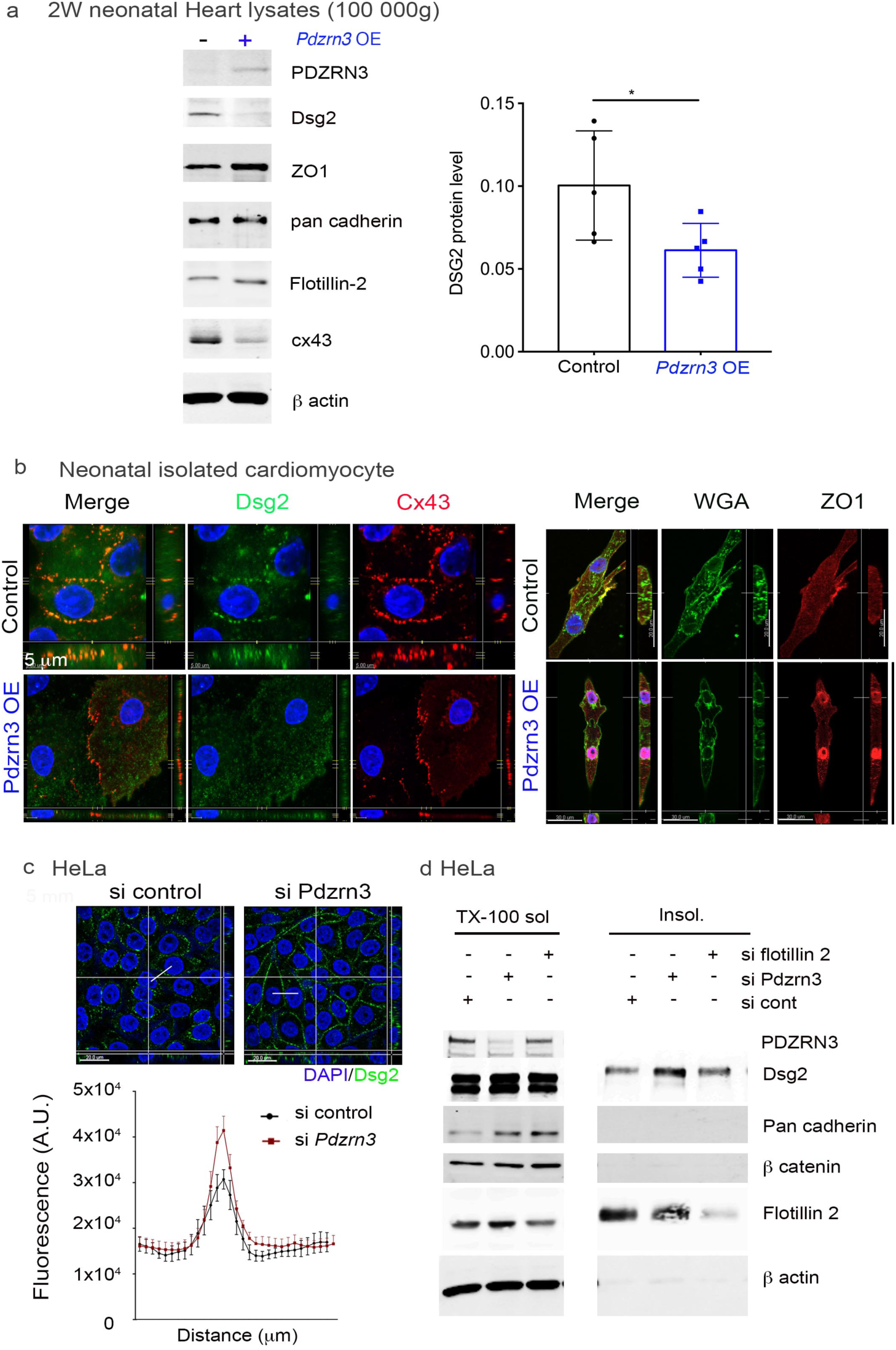
Pdzrn3 regulates Desmoglein accumulation at cell membrane. a. Western blot analysis of indicated proteins in plasma membrane fraction of heart tissues retrieved from control and *Pdzrn3* OE mice at 2 Weeks of age. Relative expression of Dsg2, from Western blots. Significance vs control *, P<0.05 ; 1 by unpaired *t*-test (n=5 mice per group). b. Representative confocal microscopy images of primary cardiomyocytes from control and *Pdzrn3* OE neonates stained with antibodies against Dsg2, Cx43, WGA-Fitc and ZO1. c. After immunostaining of HeLa cells depleted of *Pdzrn3* (si Pdzrn3) or not (si control) for Dsg2 (green), fluorescence intensity profile curve were plotted. Data are represented as mean ± sem from three independent experiments. d. Western blot analysis of indicated proteins in triton-X-100 soluble and insoluble fractions from HeLa depleted either of *Pdzrn3* (si Pdzrn3) or *flotillin-2* (si flotillin 2).

## Discussion

In the heart, polar accumulation of junction proteins at the IDs is necessary for terminal differentiation of myocytes and is required for global adaptive architecture of the heart against hypertrophic stress. Here we identify a physiologically important role for the PCP regulator PDZRN3 in these processes. We report that cardiomyocyte specific *Pdzrn3* deletion, protects the heart from the transition to heart failure, under conditions of pressure overload stress. Consistently, PDZRN3 sustained expression in cardiomyocytes during the first 2 weeks after birth, lead to abnormal heart morphogenesis, cardiac eccentric hypertrophy and early death. Forced PDZRN3 expression impairs postnatal cardiomyocyte elongation, and final geometry, disrupts IDs with a decrease of Dsg2 accumulation at cell borders which coincide with a low to undetectable Cx43 expression, displacing ZO-1 toward nuclei. We found that human cardiomyocytes from patients suffering mal-adaptive hypertrophy displayed hallmarks of PDZRN3 signaling re-induction.

We show here that *Pdzrn3* controls a genetic program essential for heart maturation and for the maintenance of overall cardiomyocyte geometry and contractile function. It has previously been reported that during neonatal heart maturation, and perinatally, ZO-1 becomes gradually restricted to the ID of the myocytes, where it recruits Cx43 from the initial circumferential plasma membrane to the ID to generate functional gap junction channels(Barker et al., 2002; Toyofuku et al., 1998). Similarly, our results in mice indicate that during the brief period after birth, neonatal cardiomyocytes evolve from a round, non-polarized immature state to an elongated, aligned and highly polarized state with the cell junction proteins Cx43 and ZO-1, localized to the ID, joining cells end to end. Forced postnatal expression of PDZRN3 impaired Cx43 induction and dynamic localization of both Cx43 and ZO-1 at the IDs, as early as 1 week after birth, a time during which Cx43 gap junction plaques form at both cell ends. We report an almost complete loss of Cx43 staining after 15 d of *Pdzrn3* OE in transgenic hearts, which could be due to Cx43 degradation through lysosomal and proteasomal ubiquitin dependent pathways(James G. Laing et al., 1997). This remodeling of gap junctions may account for cardiac conduction alterations in Pdzrn3 OE mice as previously reported(Danik et al., 2004).

Formation of desmosome precedes gap junction channel(Saffitz, 2009). Dsg2 is a major and essential cadherin of the cell-cell contact in cardiac desmosomes(Eshkind, 2002) where it helps to maintain planar coordination of cardiac myocytes. Disruption of adhesive integrity via mutations in the *DSG2*-gene are regarded to cause severe arrhythmogenic cardiomyopathy(Debus et al., 2019; Krusche et al., 2011; Maron et al., 2006). In contrast, induced deletion of Cx43 in the adult heart did not affect the spatial organization of adherens junction and desmosome in the ID(Gutstein, 2003). In agreement, we observed a loss of cardiac tissue integrity with a rupture of cardiomyocyte junctions at the ultrastructural level in mutants overexpressing *Pdzrn3*. We showed here that forced expression of *Pdzrn3* did not modify significantly overall cellular level of Dsg2 but diminished Dsg2 localization at cell membrane while repressing *Pdzrn3* was sufficient to enhance Dsg2 accumulation at cell/cell contacts. Dsg2 is dynamically localized to plasma membrane, through a microtubule dependent transport; altering this mechanism results in weakened adhesive strength (Nekrasova et al., 2011). Thus, our findings suggest that PDZRN3 signaling regulates this active process of Dsg2 recruitment at cell membrane.

Both canonical and non-canonical Wnt signalling have been implicated in the multistep process of cardiogenesis (Gessert & Kühl, 2010; Jeong et al., 2017; Mazzotta et al., 2016), however, understanding of the outcome of activating and inhibiting the Wnt pathways at specific times during the development of the heart is still not fully understood. Our data seems to favour the hypothesis that in the heart, PDZRN3 is acting in the PCP pathway. In normal adult cardiomyocytes, Wnt/Fzd is quiescent and becomes reactivated in the development of pathology such as hypertrophy (Malekar et al., 2010). The stabilization of β catenin induces hypertrophic growth (Haq et al., 2003) while β catenin cardiac deletion attenuates TAC-induced hypertrophy (Blankesteijn et al., 2008; Qu et al., 2007). It was reported that JNK overexpression, *in vivo*, led to a lethal cardiomyopathy (Petrich et al., 2004). We report that PDZRN3 overexpression, in the first 2 weeks after birth, leads to an increase of PKCζ levels with translocation of c-Jun in the nucleus, correlated with a decrease of cytosolic ß catenin without modification of the membrane ß catenin pool. This is consistent with reports showing a role of the non-canonical Wnt pathway during cardiogenesis which destabilizes ß catenin dependent canonical Wnt signals (Abdul-Ghani et al., 2011). Our data suggest that PDZRN3-induced expression of the PCP signalling factors during this critical postnatal window may contribute to the pathogenesis of heart disease.

Previous reports have demonstrated a decrease of Cx43 with a reduction in size and abundance of gap junctions in cardiac tissues retrieved from patients with ischemic, or dilated cardiomyopathy or in end-stage heart failure (Bruce et al., 2008). Moreover, ZO-1 levels were reported to be increased at the IDs in these patients, and negatively correlated with Cx43 levels at ID. Other groups have found that ZO-1 and Cx43 levels were reduced concomitant with a reduction in co-localization between these two proteins, at IDs in humans with end-stage heart failure (Kostin, 2007; J. G. Laing et al., 2007). Additionally, the lateralization of Cx43 to adherens junction-sparse regions of the membrane has been reported in tissue from the myocardial infarct border and zones of myofiber disarray in patients with hypertrophic cardiomyopathy (Sepp et al., 1996). The lateralization of Cx43 observed in many pathological scenarios is reminiscent of arrangements observed during maturational growth of the ventricle. In this pathological recapitulation of developmental patterns in the adult, cell-cell adhesion junctions appear to remain prominently located at the ID, whereas Cx43 distributes to lateral sarcolemma (Severs et al., 2008). During atrial dilation or permanent atrial fibrillation, there are a number of de-differentiated myocytes with lateralized connexins, altered ID and loss of polarity (Hong et al., 2012; Smyth et al., 2010). As for relevance of our finding, it is noteworthy that we report an increase of *Pdzrn3* levels in patients suffering from HF-HCM and coincidentally a loss of Cx43 labelling in large tissue areas along with an alteration of ZO-1 distribution compared to those suffering from HF-DCM or patient presenting and adaptive hypertrophy (control-HCM). We propose that upregulation or maintenance of *Pdzrn3* and decrease of Cx43 expression may lead to a maladaptive response to hypertrophy.

In conclusion, our data expands the current knowledge regarding early determination of cardiac tissue development and highlights a potential indication for treating the development of hypertrophic cardiomyopathies. Approaches targeting PDZRN3 signaling cascades in cardiomyocytes might be a novel therapeutic target for human heart failure.

## Materials and methods

### Experimental animals

This study was conducted in accordance with both Univ. Bordeaux institutional committee guidelines (committee CEEA50) and those in force in the European community for experimental animal use (L358-86/609/EEC).

For cardiomyocyte *Pdzrn3* overexpression (Pdzrn3 OE mice), *MHC-tTA* males (Tet-off)) (*51*) were mated with *Pdzrn3-V5* mutant females (*19*) which results in the generation of mice with genotypes of *MHC-tTA*; *Pdzrn3-V5* (Pdzrn3 OE) and Pdzrn3-V5 (control). Tails of pups were genotyped by PCR using the P1 and P2 primer set to detect the *Pdzrn3-V5* coding gene (P1, 5′-CAGCTTGAGGATAAGGCGCT-3′; P2, 5′-CTTCGAGCTGGACCGCTTC-3′) and using the P3 and P4 primer set to detect the *tTA* coding gene (P3, 5′-GCTGCTTAATGAGGTCGG-3′; P4, 5′-CTCTGCACCTTGGTGATC-3′.

For cardiomyocyte–specific deletion, with αMHC-MerCreMer transgenic mice (Sohal et al., 2001) were crossbred to *Pdzrn3* ^*f/f*^ mice (R. N. Sewduth et al., 2014), which results in the generation of mice with genotypes of αMHC-MerCreMer ; *Pdzrn3* ^*f/f*^ *(MCM-Pdzrn3*^*f/f*^*) and Pdzrn3* ^*f/f*^. For *Pdzrn3* gene deletion in adults, 0.5 mg of tamoxifen was injected intraperitoneally for three successive days, 2 weeks before surgery. Doxycycline was administrated in the drinking water (0.4 mg/ml) to inhibit cardiac specific expression of transgenic *Pdzrn3*.

Isolation of neonatal cardiomyocytes: at P3, mice were euthanized by decapitation. Hearts were carefully removed from the thoracic cavity, immediately placed into ice-cold cardioplegic solution. Ventricular neonatal cardiomyocytes were then isolated and prepared as recommended (Pierce™ primary cardiomyocyte isolation kit).

### Tissue sampling from human myocardium

Tissue biopsies from left ventricular (LV) or septal myocardium were obtained from still-beating hearts immediately after explantation from patients with end-stage heart failure (HF) undergoing heart transplantation at the Bordeaux University Hospital (NYHA class III or IV; LVEF <35%). Based on the extensive clinical record of patients, sample were classified as dilated cardiomyopathy (HF-DCM) (mainly ischemic dilated cardiomyopathy) or decompensated hypertrophic cardiomyopathy (HF-HCM). LV myocardial biopsies were obtained at the time of surgery (ventricular septal myectomy) of aortic valve replacement for severe aortic stenosis (control-HCM).

Aliquots for mRNA analyses were snap-frozen in liquid nitrogen, and stored at −80°C until use. Tissue samples were fixed in paraformaldehyde (PFA) 4%-PBS and paraffin embedded. Procurement of human myocardial tissue was performed under protocols and informed consent approved by Biologic Resource Center and the ethical committee at Bordeaux University

## Experimental protocols

### Mouse model of cardiac hypertrophy

Tamoxifen induced *Pdzrn3* ^*f/f*^*and MCM-Pdzrn3*^*f/f*^ mice, 8 weeks old, were subjected either

- Angiotensin (Ang) II by the use of subcutaneously implanted miniosmotic pump (Alzet) for 4 weeks filled with either Ang II (125 ng/kg/min) or saline solutions.
- Or Transverse aortic contriction (TAC) by tying a 6.0 nylon suture ligature against a blunt 27-gauge needle placed adjacent to the aorta as previously described (Plateforme Genotoul anexplo, Toulouse). The needle was then removed, and the chest and overlying skin were closed. Mice were monitored up to 16 weeks after TAC procedure.

### Echocardiography

Mice were anesthetized using 1.5% oxygenated isoflurane by inhalation. Echocardiography was performed using a Visualsonics Series 2100 high-resolution imaging system with a 38 MHz Microscan transducer probe. Cardiac ventricular dimensions and fractional shortening were measured in 2D mode and M-mode 3 times for the number of animals indicated.

### Electrocardiography

Electrocardiograms were recorded from mice sedated with low dose of isoflurane using the standard four limb leads. Waveforms were recorded with AD instruments Animal Bio Amps ECG and intervals were measured manually using Powerlab device and LabChart software.

### RNA preparation and quantitative PCR

Mouse tissues were homogenized in TRI-REAGENT ™ (Euromedex) and RNA was extracted according to the manufacturer’s instructions. Q-PCR was performed as described previously (*19*). The following primers sets were used: mouse cyclophyllin (NM_009505), F: 5’-AGCTAGACTTGAAGGGGAATG -3’, and R: 5’-ATTTCTTTTGACTTGCGGGC-3’; mouse *Nppa* (NM_008725), F: 5’-CGTCTTGGCCTTTTGGCTTC -3’, and R: 5’-GGTGGTCTAGCAGGTTCTTGAAA -3’; mouse *Nppb* (NM_008726), F: 5’-AAGCTGCTGGAGCTGATAAGA -3’, and R: 5’-GTTACAGCCCAAACGACTGAC -3’; mouse *Pdzrn3*, F : 5’-CTGACTCTTGTCCTGCATCGGGACTC-3’, and R: 5’-ATGGGC TCCTTGGCTGTCTTGAAAGC-3’, mouse *Gja1* (NM_010288), F: 5’-ATCAGGGAGGCAAGCCATGCTCA -3’, and R: 5’-ACGTTGGCCCACACCACAAAGA - 3’. For human tissues, GAPDH, (NM_001289746), F: 5’-GAGTCAACGGATTTGGTCGT - 3’, and R: 5’-TTGATTTTGGAGGGATCTCG -3’ *Gja1* (NM_000165), 5’-AAAGTCAGGGAATCAAGCCATGCTT -3’, and R: 5’-ACACCATATTGGCCCACACCACA-3’; human Pdzrn3, F : 5’-CAGCTTGAGGATAAGGCGCT-3’, and R: 5’-CTTCGAGCTGGACCGCTTC-3’. Target gene expression was normalized to that of the control gene, and relative expression was quantified by the comparative Ct method (2^-ΔΔCt^).

### Histology and immunohistochemistry

For tissue immunohistology analysis, hearts were fixed in paraformaldehyde (PFA) 4%-PBS, then paraffin embedded and sections were stained with haematoxylin/eosin for morphologic evaluation, picrosirius red for fibrosis quantification and labeled with tetramethyl rhodamine isotiocyanate-labeled wheat germ agglutinin (WGA, Sigma-aldrich) for cross sectional area. Captured images using a Zeiss (Axioimager) were analyzed with the Image J analysis software. Immunofluorescence staining was performed as previously described(Raj N. Sewduth et al., 2017) with antibodies specific for CD45 (BD pharmingen), Connexin 43 (Sigma), ZO1 (Invitrogen), β catenin (sigma-aldrich), N Cadherin(Santa cruz). Images were taken with a Zeiss LSM700 confocal laser-scanning microscope confocal microscope.

Transmission electron microscopy (EM) was performed on glutaraldehyde-perfused hearts as previously described (Raj N. Sewduth et al., 2017).

### FTIR analysis

Formaline fixed-hearts were then snap frozen. A 20 µm section was then deposited onto an IR-transparent ZnS window (Crystan, UK) for FTIR analysis. Four sections from the left ventricle were analyzed. For each section, about 50 IR spectra were recorded. IR spectra were collected in transmission mode using a Spotlight 300 FTIR imaging system, coupled to a Spectrum One spectrometer (Perkin-Elmer). Data analyses were carried out with the OPUS 7.2 software sub-routine (Bruker, Germany). Pre-processing of baseline correction and normalization spectra were applied. Then spectra were used for classification (hierarchical cluster analysis) using Ward’s algorithm. Classifications were performed on the second derivative of the spectra with 7-points smoothing using 1200-900 cm^-1^ spectral interval from the IR spectra. Then the average of the IR spectra from a cluster was performed and the average were compared (Hackett et al., 2013).

### F/G actin ratio measurements

The amount of F-actin and G-actin was measured with an actin polymerization assay kit (BK037, Cytoskeleton). 2 week-old mice were sacrificed by cervical dislocation. For each sample, 2 hearts were sliced into small pieces with a scalpel and quickly homogenized in 1 mL of F-actin stabilization buffer with ATP and protease inhibitorswith a Dounce homogenizer and then incubated at 37°C for 10 min. Tissue homogenates were spun at 100,000xg for 1 h at 37°C to separate the Globular (G)-actin (supernatant) and filamentous (F)-actin fractions. The pellets were resuspended to the same volume as the supernatant in using ice-cold Milli-Q water and 10 μM cytochalasine D (F-depolymerizing solution) and left on ice during 1 h. Actin was quantified by Western blot using an anto-actin antibody (Cytoskeleton). The F-actin/G-actin ratio was determined using the Odyssey infrared imaging system (LI-COR Biosciences).

### Cell culture and transfection

HeLa cells were cultured in RPMI supplemented with 10% fetal bovine serum and penicillin-streptomycin. SiRNA were transfected using Interferin (Polyplus) at a final concentration of 30nM. The oligonucleotides used were designed by Origene for h-pdzrn3 (Raj N. Sewduth et al., 2017).

For subcellular triton soluble and insoluble fractions, the cells were scraped with a plastic blade in 10 mM Tris pH 7.5, 140 mM NaCl, 5 mM EDTA, 2 mM EGTA and 1% Triton X100 supplemented with 0.1 mM sodium orthovanadate, 0.1 mM PMSF, 20 µg/ml leupeptin, benzonuclease, 50 µg/ml pepstatin, aprotinin, trypsin inhibitor and 0.1 µM okadaic acid. culture dishes then the mixture was incubated for 10 min on ice and centrifuged for 5 min at 800×*g*. The supernatant was then centrifuged for 30 min at 14 000×*g*. The supernatant was saved (Triton-X100-soluble fraction) and the pellet was dissolved in in 10 mM Tris pH 7.5, 8M uree, 140 mM NaCl, 5 mM EDTA, 2 mM EGTA and 1% SDS (triton-X100 insoluble fraction).

### Western blot, and fractionation

Immunoblotting were performed as described previously (Raj N. Sewduth et al., 2017). For tissue fractionation, tissue lysates were processed with cell extraction kits (Thermo scientific Pierce) as recommended. Briefly, cellular compartments are sequentially extracted by incubating cells with cytoplasmic, membrane and nuclear protein buffers.

Proteins were then resolved by SDS–polyacrylamide gel electrophoresis and blotted with antibodies specific for V5 (Invitrogen), PDZRN3 (Santa Cruz), ZO1 (Invitrogen), β-catenin (Sigma-Aldrich), PKCζ (Santa Cruz Biotechnology), c-Jun (Upstate), Cx43 (Sigma) α-tubulin (Sigma-Aldrich), Dsg2 (Acris Origene), pan cadherin (Sigma), Flotillin-2 (Santa cruz). Binding of antibodies to the blots was detected using the Odyssey Infrared Imaging System (LI-COR Biosciences).

For mass spectrometric analysis and protein identification, proteins recovered within V5 immunoprecipitates were excised from colloidal coomassie blue–stained gels. Peptide sample preparation and LC-MS/MS analysis was performed in collaboration with the proteome plateform of the Functional Genomic Center of Bordeaux (CGFB). Briefly, the excised gel pieces were destained, reduced with DTT and alkylated iodoacetamide. Excess reagents were washed out and Trypsin in-gel digestion were performed. Tryptic peptides were extracted from gel plugs, dried and resuspended in 2% acetronitrile with 0.05% trifluoroacetic acid (TFA). The LC-MS/MS measurements of peptide solutions were carried out on Q-Exactive mass spectrometer (Thermo Fisher Scientific). Samples were injected on a C18 precolumn (Acclaim PepMapTM) and further separated on a 75 µm ID x 15 cm nanoviper C18 reversed phase column (Acclaim PepMap RSLC) with gradient from 4% to 40% ACN in 0.1% formic acid for 120min min at a flow rate of 300 nl/min. Full MS scans were acquired in the Orbitrap mass analyzer over m/z 300ñ2000 range.

### Statistical analysis

The experimental results represent means ± sem. Each experiment was conducted at least three times. When multiple experiments using different numbers of animals were pooled for the statistical analysis, the range of number of animals was indicated in the figure legend. Comparison of continuous variables between two groups was performed using the unpaired two-sided Mann-Whitney U-test (non-parametric). Comparison of multiple groups was performed by ANOVA. Cumulative survival data was evaluated by Kaplan-Meier non parametric regression analysis and the log-rank test. All analyses were performed with appropriate software (GraphPad Prism Software). *P* < 0.05 was considered statistically significant. The statistical test is indicated for each data analysis.

## Acknowledgments

We warmly thank the Plateforme de la génomique fonctionnelle de Bordeaux for Mass spectrometry analysis and Denis Calise from the plateforme Plateforme Genotoul anexplo, Toulouse for TAC surgical protocol. We thank Sylvain Grolleau for excellent animal care and breeding. We thank the BIC platform for electron microscopy images. This work was supported by Agence National de la Recherche (ANR -16-CE17-0001-01), by the Fédération Française de Cardiologie and by INSERM funds.

## Author contributions

C.D. and T. C. conceived the study. M.P., B.J, I.F., L.C., M.H., C.D. performed the experiments. E.B., S.H. and P.D. contributed to data interpretation. M.P., C.D. and T.C. wrote the manuscript, with contributions from all the other authors.

## Competing interest statement

The authors declare no competing financial interests

---

**Suppl. Fig. 1.**
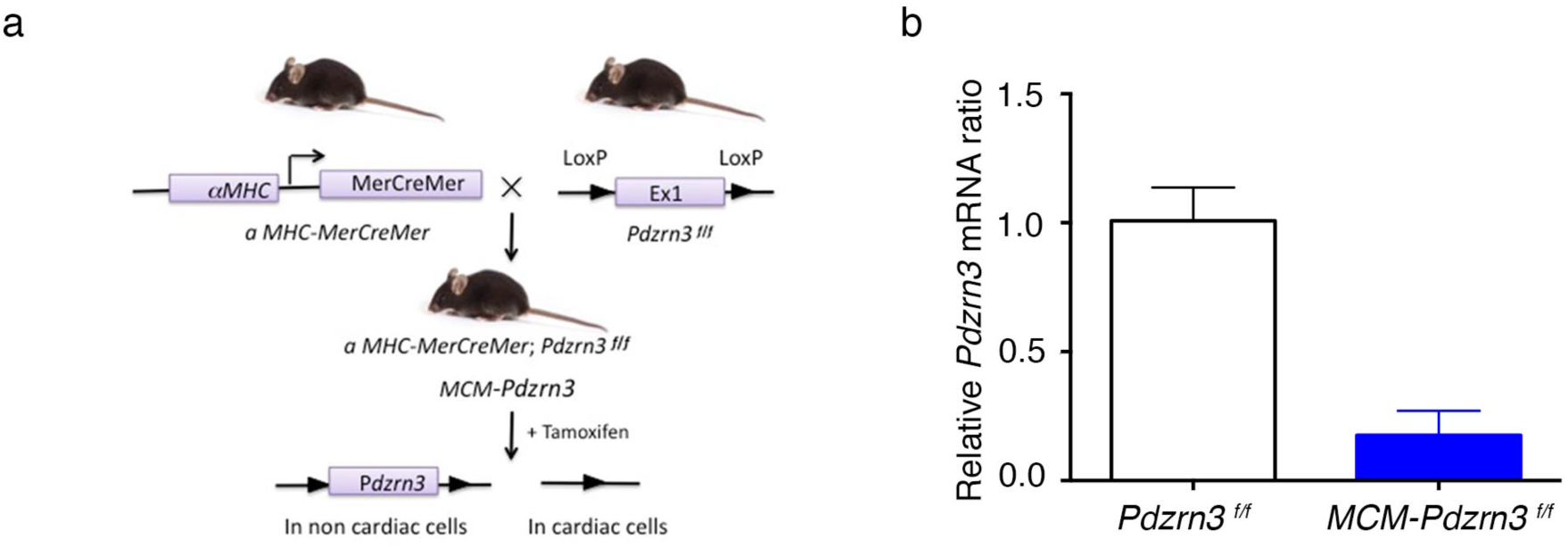
Transgenic mouse model for *Pdzrn3* depletion in cardiomyocytes: a. Schematic of *Pdzrn3* deletion in cardiac cells. Control mice were *Pdzrn3*^*f/f*^. MCM-*Pdzrn3* KO mice were MHC-MerCreMer; *Pdzrn3*^*f/f*^. b. qRT-PCR demonstrates deletion o*f Pdzrn3* in adult hearts after 1 month of tamoxifen treatment (n=5). Data are reported as average ± s.e.m. *P<0.05, by unpaired *t*-test

**Figure S2.**
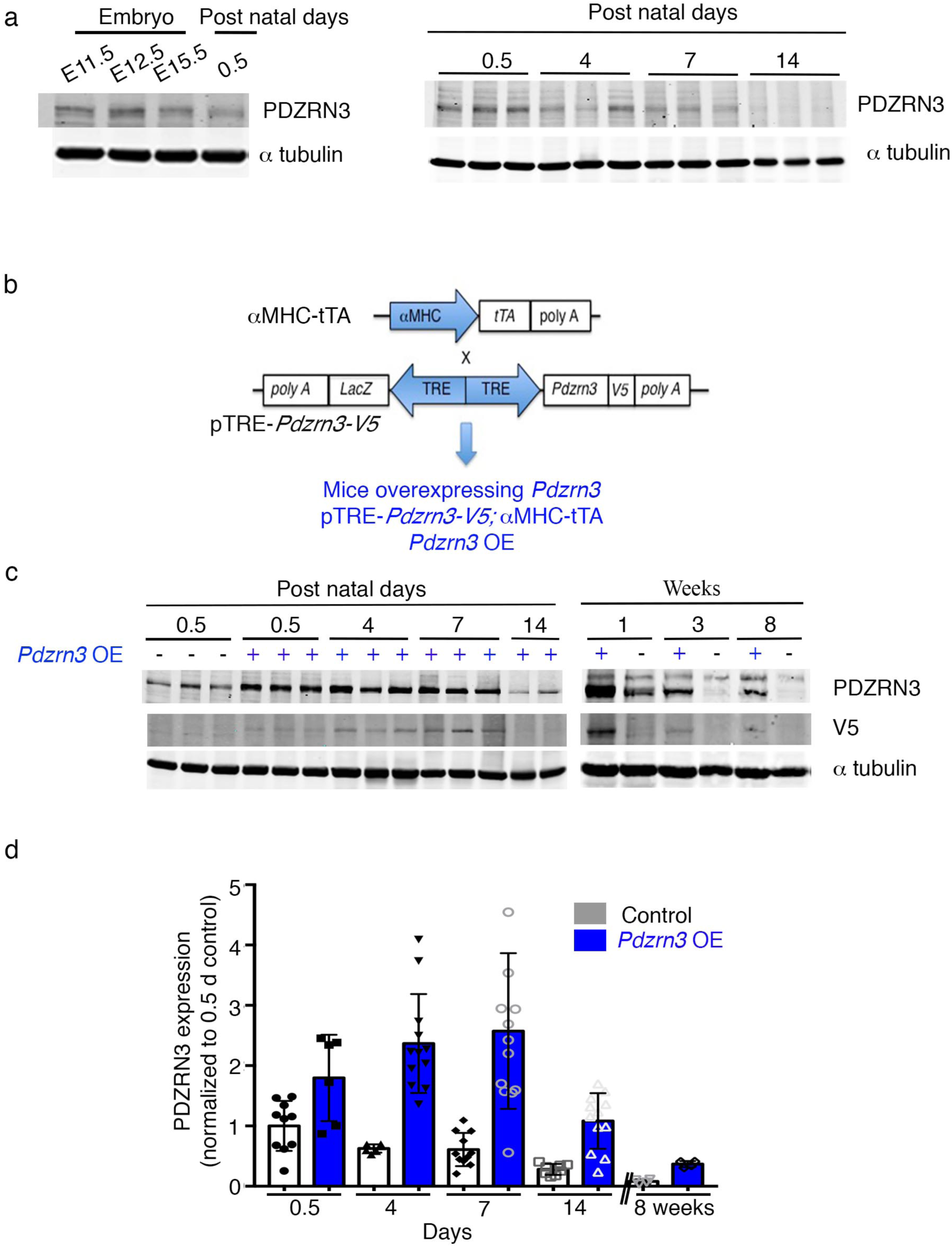
Myocardial expression pattern of *Pdzrn3* in control and *Pdzrn3* OE mice. a. Representative Western blot demonstrates levels of PDZRN3 in the embryo at E11.5, 12.5 and 15.5 days; and after birth (0.5, 4, 7 and 14 d). *α* tubulin was used as the loading control. b. Schematic of *Pdzrn3* overexpression (OE) in the heart. Control mice were *α*MHC-tTA, *Pdzrn3* OE mice were pTRE-*Pdzrn3*; *α*MHC-tTA. c. Representative Western blot demonstrates levels of PDZRN3 overexpression after birth at the indicated time points. *α* tubulin was used as the loading control. d. Representative Western blot demonstrates levels of endogenous PDZRN as well as overexpression from 0.5 to 14 days and at 8 weeks after birth. *α* tubulin was used as the loading control. The ratio is quantified and calibrated to the average of 0.5 d control mice. Data are expressed as mean ± s.e.m. (n=6-13).

**Supl Fig. S3.**
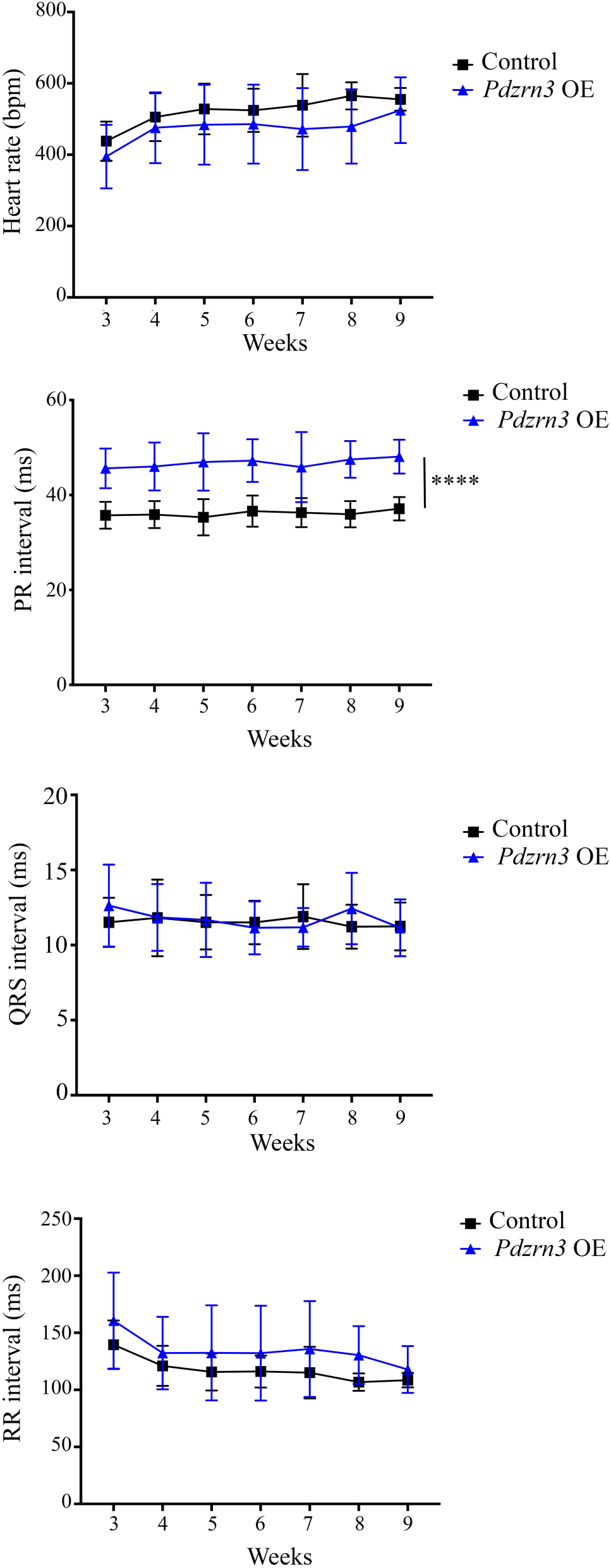
Analysis of heart rate in control and Pdzrn3 OE mice. Variation of heart rate (beat per min), as well as PR, QRS, and RR intervals (ms) in control and Pdzrn3 OE mice from 3 to 9 weeks after birth. (respectively n=6 mice vs n=4 mice) ***, P<0.001 by repeated-measures two way ANOVA with tukey’s test.

**Suppl Fig S4.**
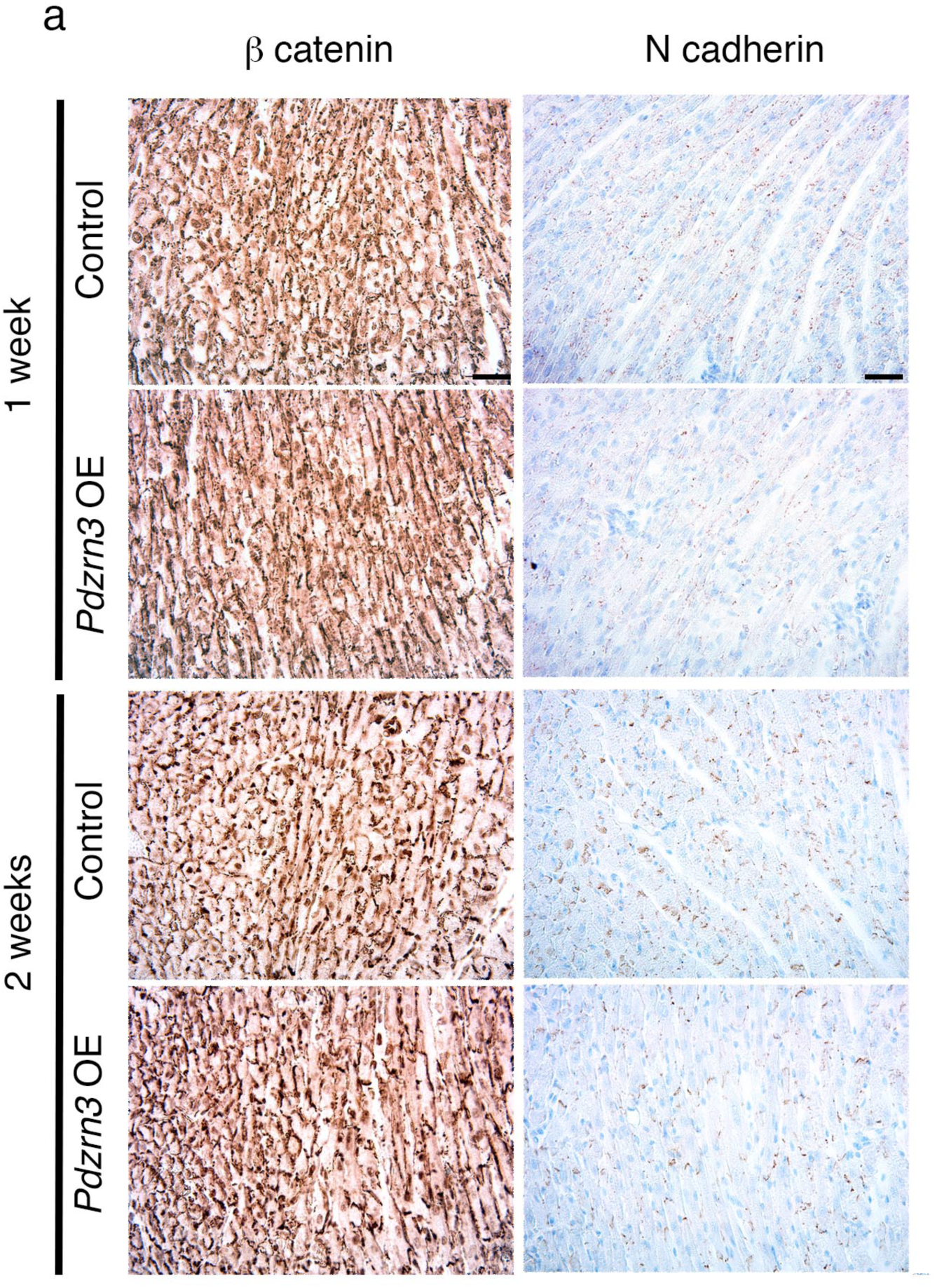
Immunolabeling of β catenin and N cadherin in control and *Pdzrn3* OE hearts at 1 and 2 weeks of age. Scale bars represent 50 μm.

**Suppl Table 1:**
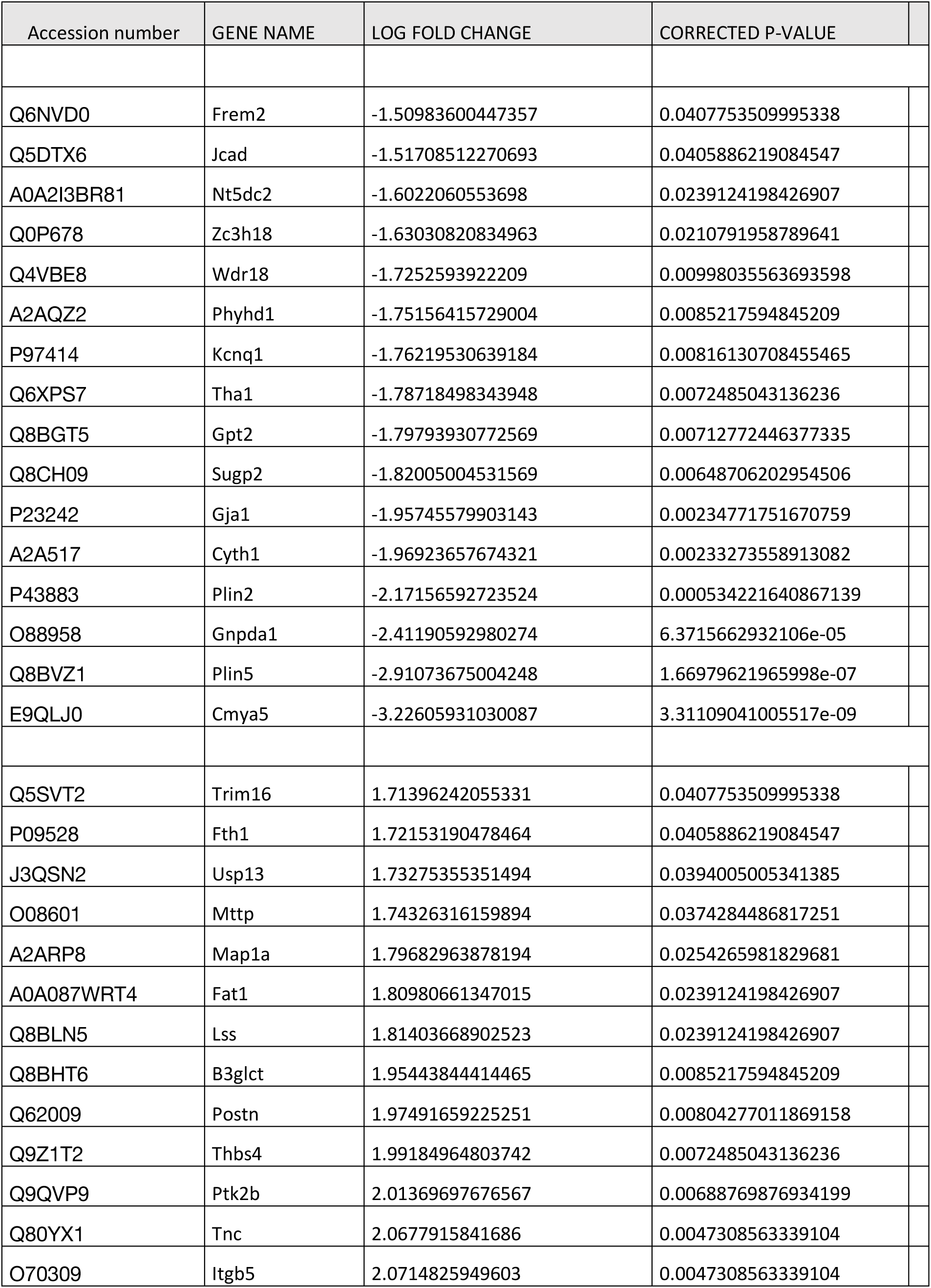
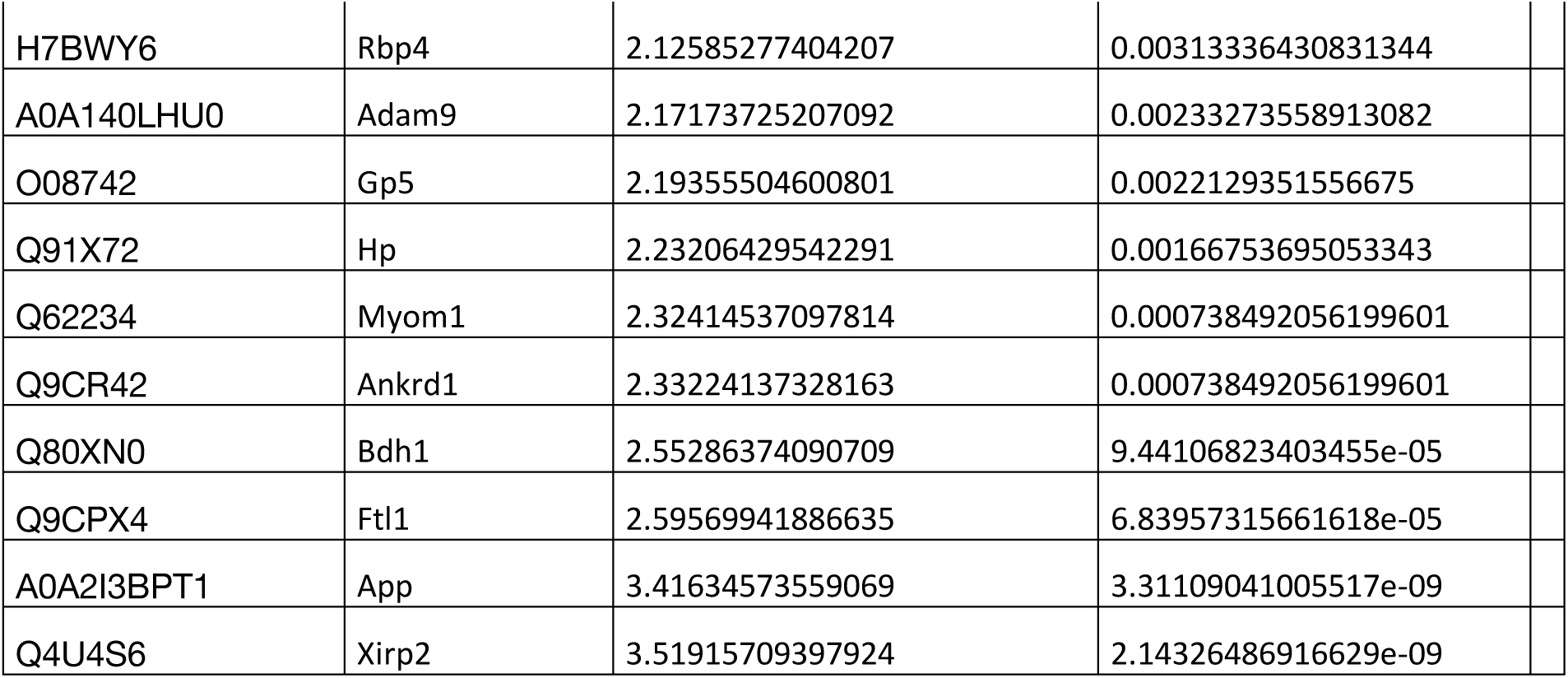
List of biomarkers down and up regulated at 14d in heart lysate from Pdzrn3 OE mice versus littermate mice

**Supplementary Table 2.**
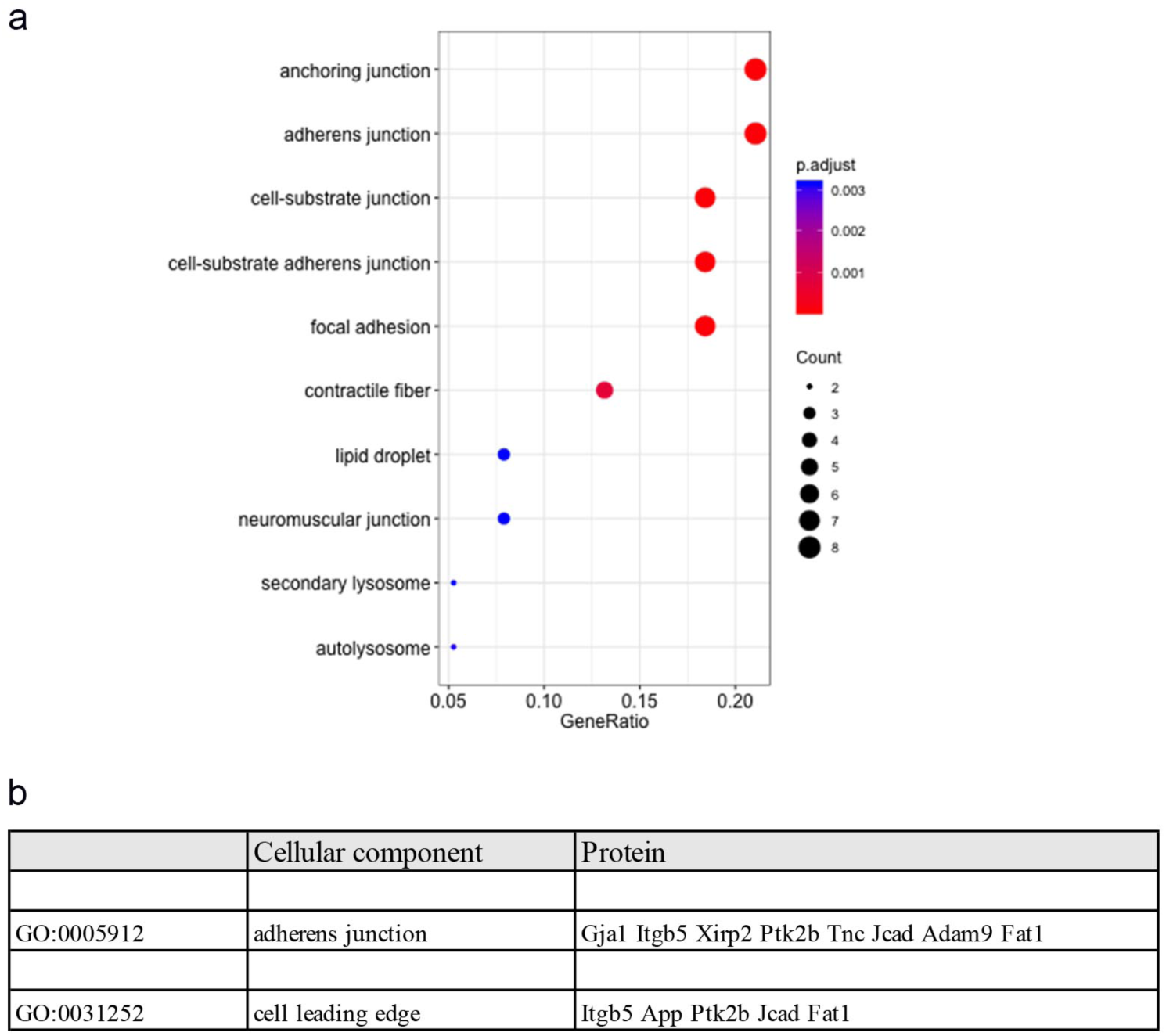
Sub-cellular functional classification of the identified proteins. a. List of identified biomarkers in adherens junction and cell leading edge pathways. b. hierarchical comparison of proteins grouped according to their similar sub cellular locations and expressed as number of assigned proteins (P<0.05)

## Notes

### Competing Interest Statement

The authors have declared no competing interest.

